# Optogenetic rescue of a developmental patterning mutant

**DOI:** 10.1101/776120

**Authors:** Heath E. Johnson, Stanislav Y. Shvartsman, Jared E. Toettcher

## Abstract

Animal embryos are partitioned into spatial domains by complex patterns of signaling and gene expression, yet it is still largely unknown what features of a developmental signal are essential. Part of the challenge arises because it has been impossible to “paint” arbitrary signaling patterns on an embryo to test their sufficiency. Here we demonstrate exactly this capability by combining optogenetic control of Ras/Erk signaling with the genetic loss of terminal signaling in early *Drosophila* embryos. Simple all-or-none light inputs at the embryonic termini were able to completely rescue normal development, generating viable larvae and fertile adults from this otherwise-lethal genetic mutant. Systematically varying illumination parameters further revealed that at least three distinct developmental programs are triggered by different cumulative doses of Erk. These results open the door to the targeted design of complex morphogenetic outcomes as well as the ability to correct patterning errors that underlie developmental defects.

**Summary:** Spatial gradients of protein activity pattern all developing embryos. But which features of these patterns are essential and which are dispensable for development? Answering this question has been deceptively difficult, due to the difficulty of shaping spatial gradients using chemical or genetic perturbations. Johnson and colleagues use optogenetics to fully restore normal embryo development with a pattern of light, opening the door to complete control over spatial organization in developing and engineered tissues.

## Introduction

During animal development the embryo is patterned by gradients of protein activity that define cells’ positions along the body axes and within developing tissues (1). In recent years, many developmental patterns have been characterized in precise quantitative detail in individual embryos (e.g. Bicoid; Sonic Hedgehog; Erk) (2–4). Yet it remains unclear which features of these dynamic, spatially-variable signaling patterns are essential (5), or how many distinct activity levels (e.g. protein concentrations or cumulative doses) of a single signaling pattern are “read out” by the genetic networks that serve as signal interpretation systems.

We can envision an idealized experiment to define the essential information contained in a developmental pattern (**Figure 1A**). First, one might prepare a “blank canvas” embryo in which a specific developmental signaling pattern is completely eliminated. On this background one might then “paint” a wide range of precisely defined candidate input patterns and then monitor their capability to rescue each developmental process. The recent development of optogenetic control over cell signaling opens the door to exactly this capability. In principle, an appropriately-tailored light input could be used to produce any spatiotemporal signaling pattern, enabling a developmental biologist to test for the essential features required to support proper tissue patterning, or allowing a bioengineer to apply non-natural stimuli to implement novel tissue architectures or morphogenetic programs (6, 7). However, despite the promise of this approach, it is still unknown whether the engineered light-sensitive proteins that make up optogenetic signaling systems are able to recapitulate all of the functions of natural developmental signals, and therefore whether a spatial light stimulus might even be capable of fully rescuing the loss of an endogenous signaling pattern.

**Figure 1.**
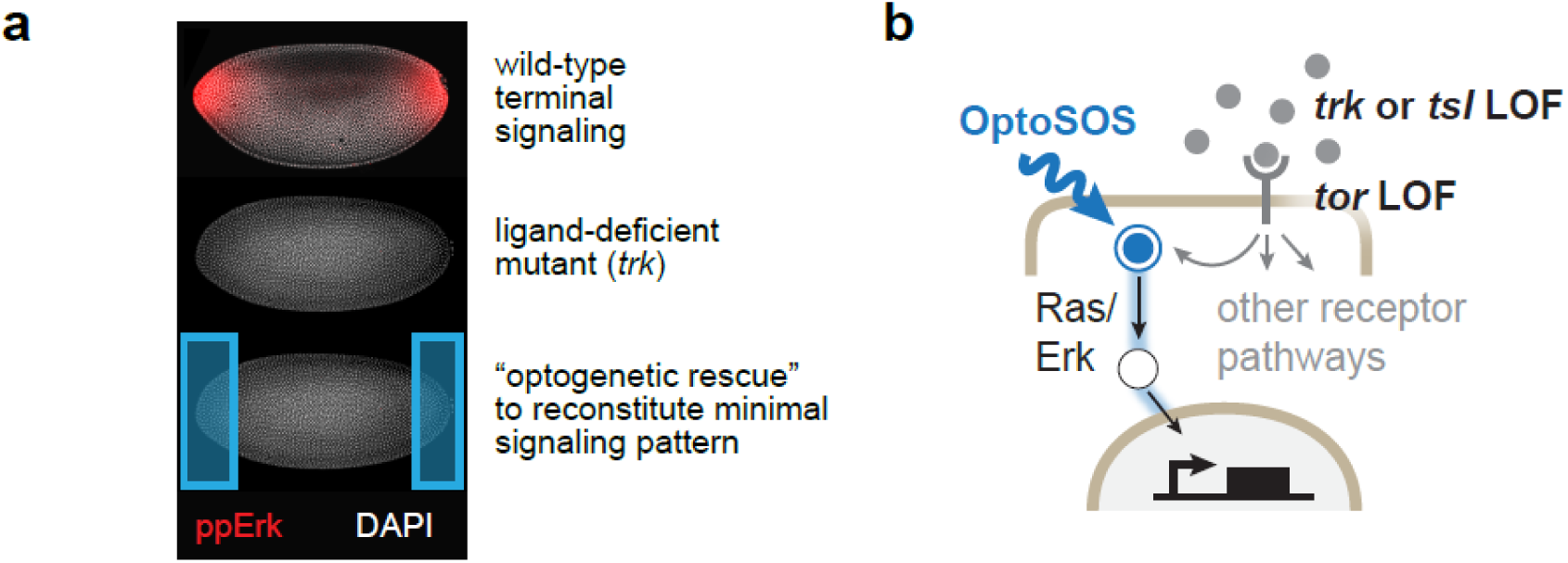
Painting developmental patterns on a blank canvas. (**A**) Upper: immunofluorescence (IF)s for doubly phosphorylated Erk (ppErk; red) in a nuclear cycle 14 embryo, exhibiting the characteristic terminal gradient. Middle: IF for ppErk in a *trk*^1^ mutant embryo, showing complete loss of terminal ppErk. Lower: Schematic of the proposed experiment, where light is applied on the *trk* mutant background to potentially restore Erk activity and function. (**B**) Receptor-level activation can be bypassed by loss of the upstream components Tor, Tsl and Trk. The light-activated OptoSOS system can be used to directly activate the Ras/Erk pathway in a loss-of-function embryo.

As a first test of this approach, we set out to perform an optogenetic rescue of terminal signaling, the first essential developmental pattern of receptor tyrosine kinase (RTK) activity during *Drosophila* embryogenesis. The terminal pattern is orchestrated by localized activation of the RTK Torso (Tor) by its ligand Trunk (Trk) at the embryonic anterior and posterior poles, leading to a reproducible terminal-to-interior gradient of Erk kinase activity that is dynamically established over a 2-hour window in early embryogenesis (4) (**Figure 1A**, top). This localized pattern of activation is essential: embryos from mothers lacking Tor, Trk, or the required co-factor Torso-like (Tsl) lack the terminal Erk gradient (**Figure 1A**, middle) and are deficient in a wide variety of anterior- and posterior-localized processes, including the formation of mouth parts and tail structures, the differentiation of many endoderm-derived tissues, and the ability to coordinate tissue movements during gastrulation (8, 9).

Here, we report the construction of OptoSOS-*trk* embryos that completely lack receptor-level terminal signaling but whose Erk activity can be controlled with light (**Figure 1B**). Remarkably, illuminating the embryo’s poles is sufficient to rescue the entire fly life cycle from this otherwise-lethal genetic mutant: illuminated OptoSOS-*trk* embryos develop normal head and tail structures, gastrulate normally, hatch, metamorphose, mate and lay eggs. We define the lower essential limits of terminal Erk signaling, demonstrating that at least three distinct developmental switches are triggered at successively increasing Erk activity doses. Our study thus demonstrates that Ras activation by SOS is sufficient to recapitulate all the essential features of receptor tyrosine kinase signaling at the embryonic termini. It also reveals that the spatial gradients of Erk activity normally observed at the termini are dispensable, as simple, all-or-none light patterns were able to rescue. Together, our data provides a first step towards defining the essential information contained in developmental signaling patterns and opens the door to optically programming cell fates and tissue movements with high precision in developing tissues.

## Results

### Establishing optogenetic Ras as the sole source of terminal pathway activity

We first set out to establish a genetic background where light could be used as the sole source of Erk activity at the embryonic termini, so that its ability to rescue subsequent development could be assessed. Terminal signaling an ideal system for such an optogenetic rescue, as all three components of its receptor-ligand system (Trk, Tor, and Tsl) are maternal-effect genes, with no essential zygotic requirements (8). Thus, in principle, one may be able to rescue the organism’s entire life cycle by reconstituting a single developmental pattern. We reasoned that loss of a receptor-level component could be combined with our OptoSOS system to control the Ras/Erk pathway (10–13), one of the major downstream effector pathways of terminal signaling. A crucial attribute of this optogenetic system is that it directly activates Ras, so OptoSOS-based activation could be combined with the genetic loss of Torso receptor activity to place Ras/Erk signaling solely under optogenetic control (**Figure 1B**).

We established embryos of three genotypes that each harbor our OptoSOS system and exhibit diminished or absent receptor-level activity: embryos produced by mothers homozygous for the *trk*^1^ amorphic allele or *tsl*^691^ hypomorphic allele, or embryos which carried a Tor RNAi construct (**Supplementary Methods**). We then monitored tissue movements during gastrulation in the presence and absence of blue light to assess whether optogenetic stimulation could compensate for the loss of terminal signaling in each case (**Figure S1**). Terminal signaling is required for invagination of the posterior midgut during gastrulation, and we previously found that illuminating OptoSOS embryos expands the domain of contractile midgut tissue across more than 80% of the embryo (12). All three alleles failed to undergo posterior midgut invagination in the dark, indicative of strong loss-of-function of terminal signaling (**Figure S1**). Blue light illumination restored tissue invagination in OptoSOS-*trk* and OptoSOS-Tor^RNAi^ embryos, but not OptoSOS-*tsl* embryos (**Figure S1**), consistent with a recent report suggesting that Tsl plays an additional, terminal signaling-independent role in gastrulation (14). These preliminary experiments suggested that OptoSOS-*trk* and OptoSOS-Tor^RNAi^ embryos are able to transition complete loss of terminal signaling in the dark to a strong gain-of-function phenotype upon illumination. However, to avoid potential concerns of incomplete knockdown or off-target effects due to the Torso RNAi construct, we chose to focus on OptoSOS-*trk* embryos for all subsequent experiments.

A successful optogenetic rescue also requires that light can be used to precisely define the domain where Erk is activated. To verify that precise spatial patterns can be achieved, we quantified the extent of SOS membrane translocation and Erk activity from a localized site of blue light illumination, and compared this spatial distribution to the natural endogenous Erk signaling gradient in nuclear cycle 14 embryos. We made use of miniCic, the first functional Erk biosensor for *Drosophila* (15), which is exported from the nucleus after phosphorylation by Erk (**Figure 2A**). We found that the intensity of OptoSOS membrane translocation and miniCic activity dropped off sharply compared to the endogenous terminal gradient, reaching baseline within 60 μm (vs > 200 μm for the endogenous pattern) (**Figure 2B-C**). Precise, local control of the Ras/Erk pathway is thus achievable using patterned optogenetic stimulation.

**Figure 2.**
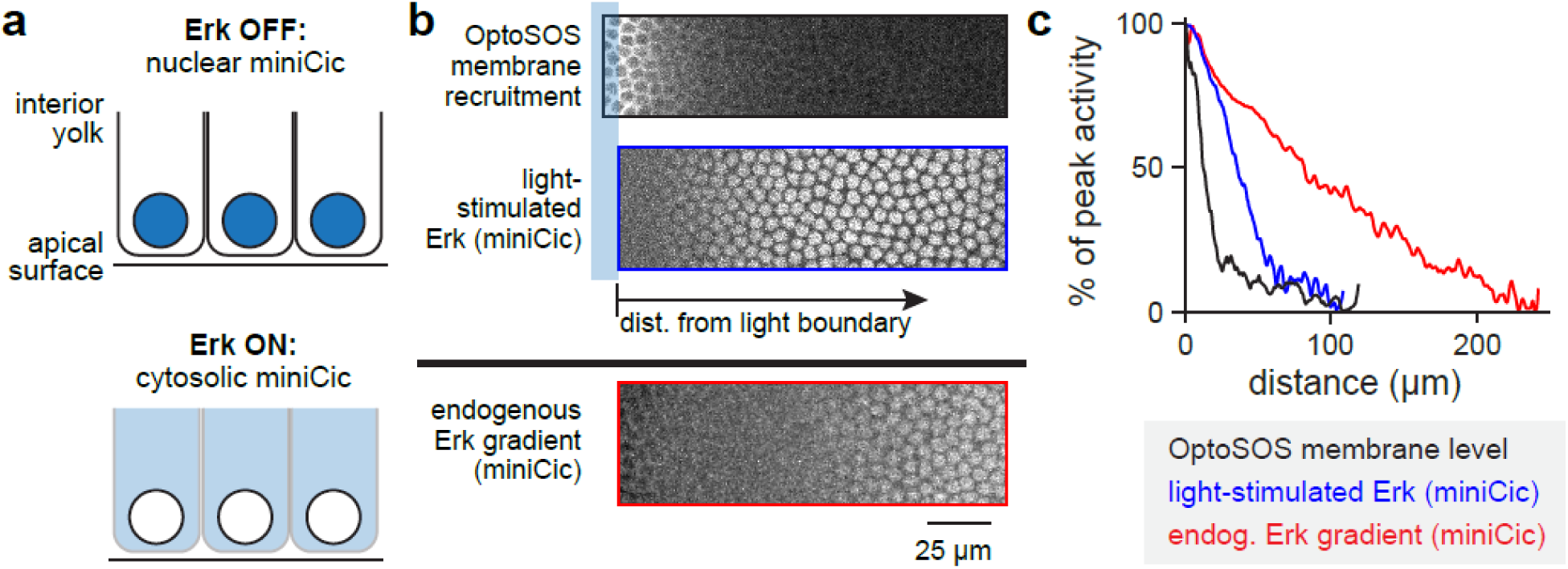
Precise spatial patterning of Erk activity using light. (**A**) A live-cell fluorescent biosensor of Erk activity (miniCic) was used to quantify the spatial distribution of Erk activity in living embryos. The miniCic biosensor is localized to the nucleus in the absence of Erk activity (left panel) and is phosphorylated and exported from the nucleus upon Erk activation (right panel). (**B**) Comparison of an all-or-none light stimulus to the endogenous terminal pattern. Regions of a representative nuclear cycle 14 OptoSOS embryo are shown, indicating OptoSOS membrane localization (top image), miniCic fluorescent intensity at the edge of an illumination pattern (middle image) or near the embryonic anterior pole (lower image). (**C**) Nuclear intensity was quantified from images in **B** as a function of position from the embryo pole (red curve) or the edge of an all-or-none light pattern (blue and black curves).

### Local illumination rescues normal development in OptoSOS-*trk* embryos

We next set out to determine whether light was sufficient for restoring various terminal signaling-associated embryonic structures in OptoSOS-*trk* embryos, and if so, which features of the light pattern might prove to be essential. We began with a simple light pattern: binary, all-or-none illumination of the anterior or posterior pole. We matched the pattern’s duration (90 min), spatial range (15% embryo length) and light intensity to roughly match the parameters observed for doubly-phosphorylated Erk during endogenous terminal signaling (**Figure S2**) (4). Although such a stimulus eliminates both the complex temporal dynamics and spatial gradient of the endogenous terminal pattern, we found that even this simple all-or-none pattern was sufficient to restore head structures that were indistinguishable from those in wild-type embryos when light was applied to the anterior pole (**Figure 3A**). Similar results were obtained upon posterior illumination, which was sufficient to restore the formation of tail structures such as posterior spiracles as well as the 8^th^ abdominal segment (**Figure 3B**).

**Figure 3.**
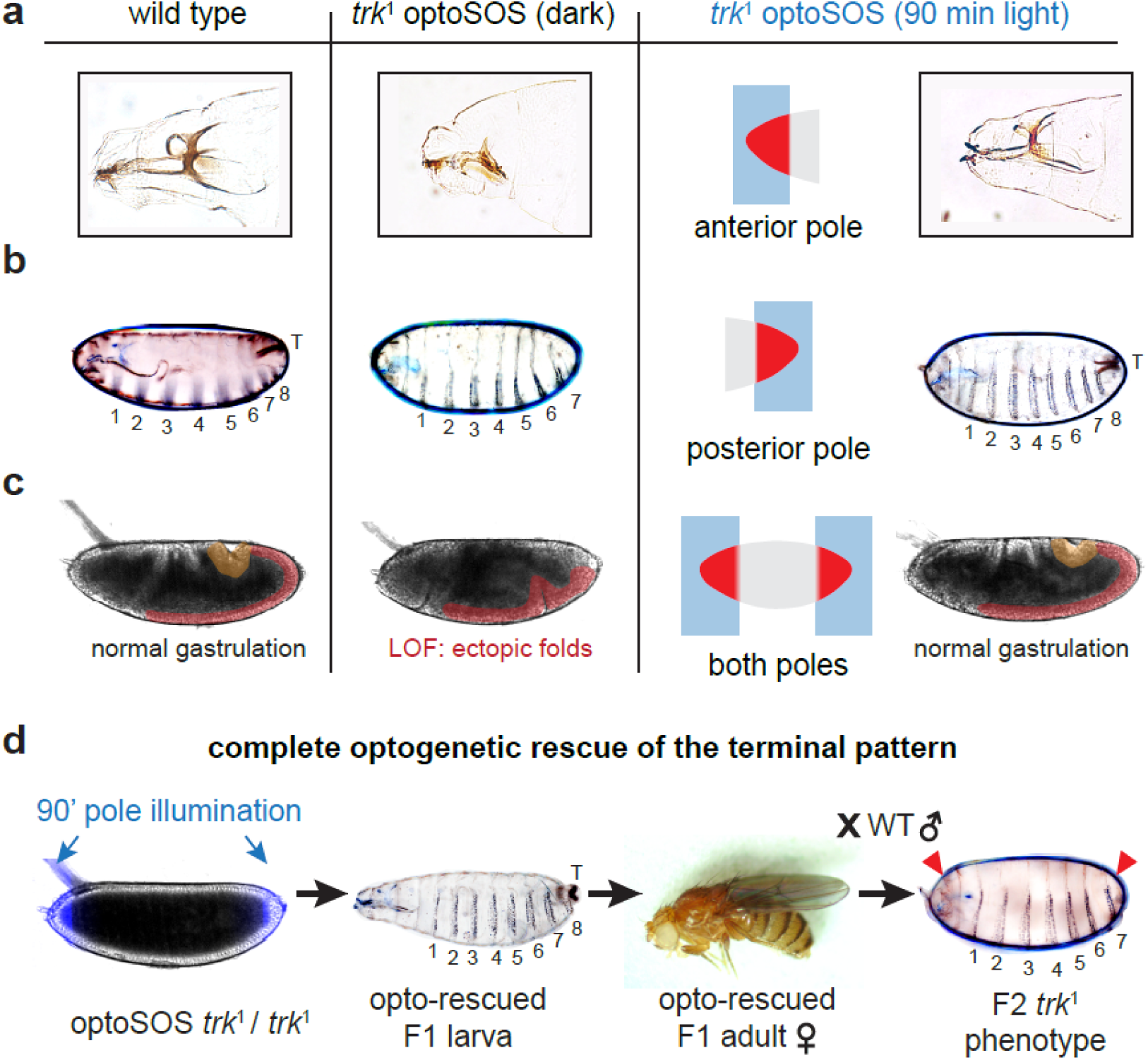
Light-controlled terminal signaling rescues normal development. (**A**) Head structures are formed normally in wild-type embryos but are absent or truncated in embryos lacking terminal signaling. Anterior illumination rescues wild-type head structures. (**B**) Tail structures (marked as “T”) and the 8th abdominal segment, missing in the absence of terminal signaling, are rescued by posterior illumination. (**C**) Rescue of normal morphogenetic movements during gastrulation, posterior midgut invagination (yellow highlight) and germ band elongation (red highlight) by illumination at both poles. (**D**) Illuminating embryos at the termini with 90 min of blue light rescues the entire fly life cycle. Embryos hatch, eclose, and mate. The embryos produced by female light-rescued flies show the *trk* mutant phenotype, as expected from this maternal-effect gene.

To assess rescue of many other terminal signaling-dependent fates and processes that are difficult to monitor individually, we next applied similar all-or-none light patterns at both poles of individual embryos and visualized the remainder of their development by differential interference contrast (DIC) microscopy. Remarkably, a majority of embryos illuminated in this manner were able to gastrulate normally, complete the remainder of embryogenesis, and hatch from the imaging device (**Figure 3C**; **Movie S1**). The larvae that hatched on the microscope were collected and maintained in standard tubes, where they proceeded normally through each instar, pupated and produced normal adult flies (**Figure 3D**). We reasoned that optogenetically-rescued female adult flies produced in this manner should still be *trk*-null, so embryos produced by these females should still harbor phenotypes consistent with the loss of terminal signaling (e.g., head defects, absence of the 8^th^ abdominal segment and tail structures). Indeed, 100% of embryos laid from light-rescued mothers failed to hatch, and cuticle preparations revealed the *trk* phenotype in all progeny (**Figure 3D**). Taken together, these data confirm the optogenetic rescue of terminal signaling in *Drosophila* embryogenesis. Simple illumination patterns were sufficient to overcome lethal defects in body segmentation, tissue morphogenesis, and cell differentiation to restore the entirety of the fly’s life cycle.

### At least three doses of terminal signaling trigger distinct developmental programs

The rescue of all anterior and posterior tissue responses by a single light dose is consistent with two different models of terminal cell fate choice. First, all terminal fates may be triggered by a single ‘decoding module’, activating all cellular responses at a single threshold of pathway activation (16). Alternatively, individual fates may be rescued one by one as the signaling dose crosses distinct fate-specific thresholds (17). To discriminate between these possibilities, we set out to map which terminal fates were triggered as the light dose was progressively increased. We found that even the lowest light dose tested was sufficient to restore tail structures in a majority of embryos (**Figure 4A**; **Figure S3**). A 5 min light pulse, delivered globally to the entire embryo, was sufficient to rescue tail structures at their proper embryonic position, revealing that these structures are triggered by a developmental switch that does not require any local spatial information from the terminal pattern. Other terminal phenotypes, including abdominal segmentation and posterior midgut invagination, were unaffected by the 5 minute Erk pulse, and still exhibited their terminal loss-of-function phenotype.

**Figure 4:**
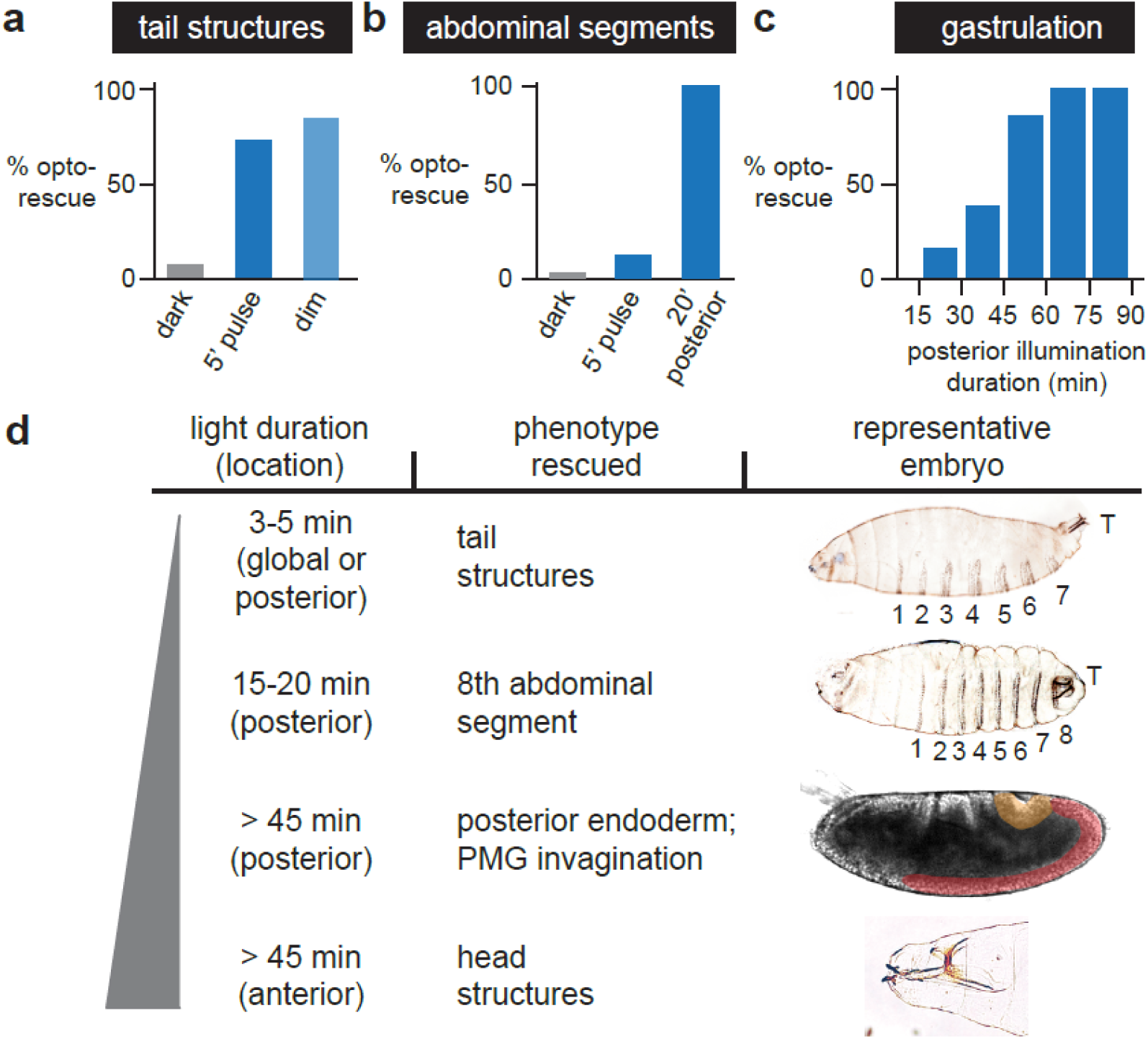
Three doses of terminal signaling trigger distinct developmental programs. (**A**) The fraction of embryos with normal tail structures was quantified from cuticle preparations from embryos incubated in the dark, subjected to a 5-minute light pulse (“5’ pulse”), or subjected to continuous, low-intensity Ras/Erk signaling driven by 1 sec blue light pulses delivered every 120 sec (“dim”). (**B**) The fraction of embryos with 8 abdominal segments was quantified from cuticle preparations from embryos incubated in the dark, subjected to a 5-minute light pulse (“5’ pulse”) or illuminated for 20 min at the posterior pole. (**C**) Posterior tissue movements during gastrulation (yellow and red highlights) were scored by differential interference contrast (DIC) imaging of individual embryos illuminated at the posterior pole for different durations. (**D**) Developmental sequence of terminal phenotypes rescued at progressively higher doses of signaling. In each case, the dose, spatial position, developmental phenotype and representative image of a rescued embryo are shown (representative OptoSOS-*trk* gastrulation and head structure images reproduced from Figure 3A and 3C).

As we progressively increasing the light dose, we observed the sequential rescue of additional developmental processes. The 8^th^ abdominal segment was restored as the posterior light stimulus was increased to 20 min (**Figure 4B**), and normal gastrulation movements were restored at 45 min of posterior illumination (**Figure 4C**). A similar dose was also required at the anterior pole for the formation of head structures. We thus conclude that Ras/Erk activity is interpreted into at least three all-or-none developmental programs with thresholds spanning nearly an order of magnitude (5 min – 45 min) (**Figure 4D**). Our data are thus consistent with a ‘multiple-threshold’ model of terminal signal interpretation: different doses of terminal signaling are able to trigger qualitatively-different downstream response programs (17). Importantly, the multiple-threshold model is perfectly compatible with our prior observation of optogenetic rescue by a single, high dose of light. That is because phenotypes appear to be rescued independently from one another, so a given light dose rescues all developmental processes that are triggered at thresholds at or below this level.

### Gastrulation movements are robust to variation in the spatial range of terminal patterning

The preceding experiments define the temporal requirements for terminal signaling, but what rules govern its permissible spatial parameters? We can again envision two extreme models. First, it is possible that only a tight range of spatial pattern widths can support normal development, by balancing the proportion of cells committed to terminal and non-terminal fates. At the other extreme, many different spatial patterns (e.g. both broad and narrow illumination widths at the termini) could funnel into a proper developmental outcome (18), resembling the tolerance to variation in the Bicoid morphogen gradient as gene dosage is varied (19) or the Shh gradient in the neural tube of Gli3^-/-^ mice (20).

To test these possibilities, we next set out to map how the spatial domain of terminal signaling affects one model developmental response: tissue morphogenesis during gastrulation. Prior to gastrulation, terminal signaling triggers differentiation of posterior endoderm tissue. This tissue then moves across the embryo’s dorsal surface during gastrulation and germ band elongation, a process that is thought to be driven by a combination of ‘pushing’ by elongating ventral tissue (**Figure 5A**; red) and ‘pulling’ by invagination of the posterior endoderm itself (**Figure 5A**; yellow) (21, 22). Terminal signaling is absolutely required, as embryos derived from *trk*-mutant mothers completely fail to undergo posterior invagination and germ band elongation, leading to buckling of the elongating tissue along the embryo’s ventral surface (8).

**Figure 5.**
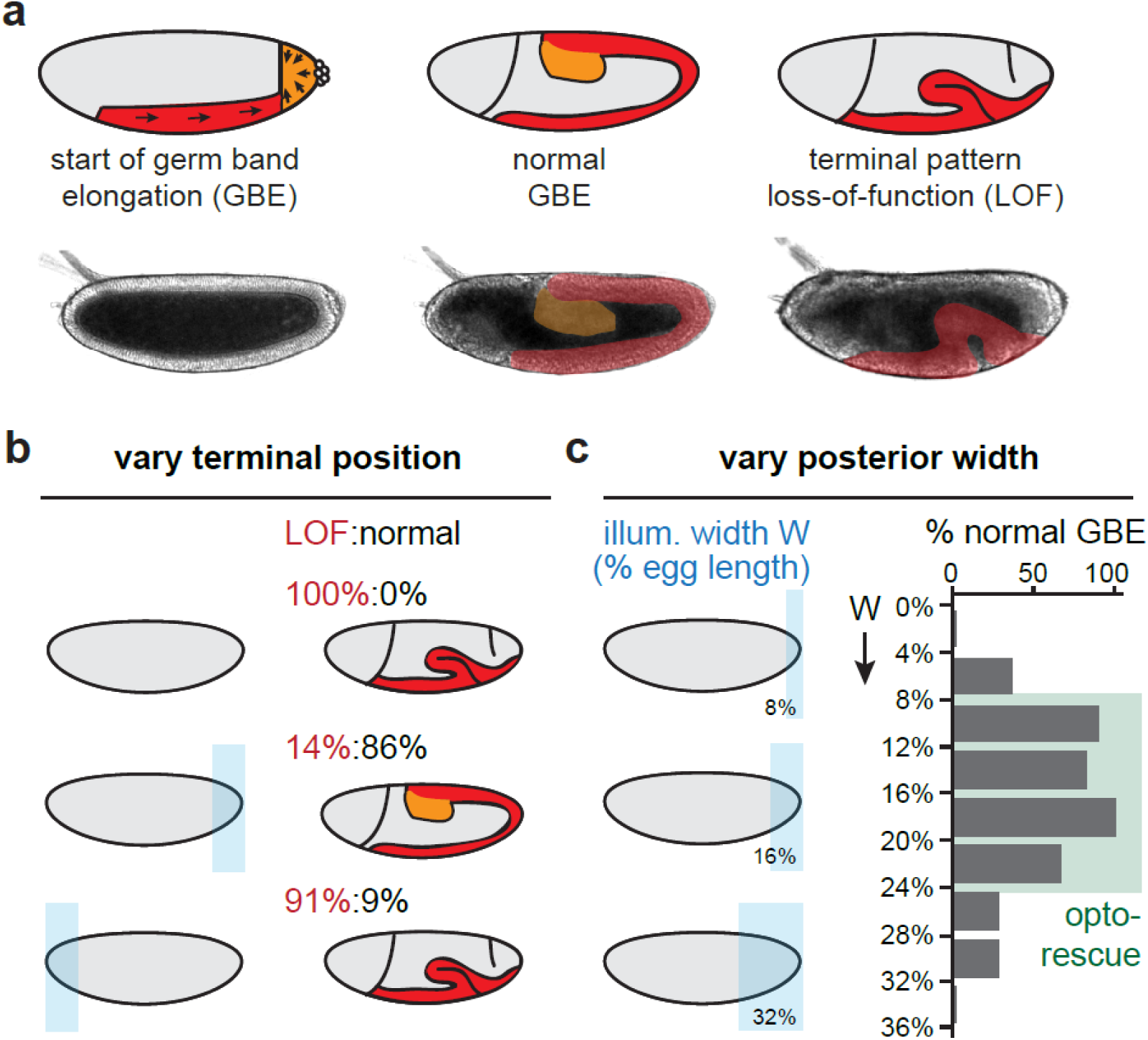
The lower limit of spatial signaling required for tissue morphogenesis. (**A**) Schematic of posterior tissue movements during germ band elongation (GBE), with representative images for each phenotype shown below. Left: During germ band elongation, cells migrate to the ventral surface (red highlight) and intercalate, while the posterior endoderm (orange highlight) constricts and internalizes. Middle: This results in the large-scale movement of posterior tissue along the dorsal surface towards the head. Right: In the absence of terminal signaling, germ band elongation is blocked, leading to buckling of ventral tissue. (**B-C**) Optogenetic dissection of the spatial requirements for terminal signaling in gastrulation. “Normal” gastrulation was defined as successful posterior invagination and a germ band whose length was within the 95% confidence interval obtained from 50 wild-type embryos. A loss-of-function (“LOF”) embryo was defined by a lack of posterior invagination and germ band elongation. (**B**) Gastrulation was scored for embryos that were kept in the dark or illuminated at either the anterior or posterior pole and monitored by differential interference contrast (DIC) imaging. (**C**) Gastrulation was scored for embryos that were illuminated with different posterior pattern widths, from 0-36% of the embryo’s length. Illumination widths that produce normal gastrulation in a majority of embryos are marked with a green box. For **B-C**, normal gastrulation is defined by presence of posterior invagination and a germ band that extends to a length within the 95% confidence interval measured across 27 wild-type embryos.

To assess whether the anterior or posterior terminal pattern is sufficient on its own for normal gastrulation, we monitored morphogenetic movements in individual OptoSOS-*trk* embryos illuminated at either the anterior or posterior pole (**Figure 5B**). We found that gastrulation failed in dark-incubated embryos as well as > 90% of embryos illuminated only at the anterior pole. In contrast, posterior pole illumination triggered normal morphogenetic movements that were quantitatively indistinguishable from wild-type embryos or embryos illuminated at both poles. Having thus determined that posterior terminal signaling is sufficient for gastrulation, we proceeded to systematically vary the width of posterior pattern. We found that the overall size of the posterior invagination scaled in proportion to the illumination width (**Figure S4A-B**), confirming that OptoSOS stimulation is indeed sufficient to program posterior endoderm (13). Yet despite the different proportion of terminal vs non-terminal tissue, a majority of embryos underwent germ band elongation that was quantitatively indistinguishable from wild-type controls as the light pattern was varied from 8–24% of the egg’s length (**Figure 5C**; **Figure S4C**). These data reveal a surprising degree of robustness in tissue morphogenesis. As the relative population is shifted between two mechanically-active cell populations (i.e., elongating ventral cells and constricting posterior cells), the speed and extent of tissue movement is unaffected. The ability to program tissue movements at any spatial positions of interest will likely make OptoSOS-*trk* embryos a rich resource for informing and challenging models of tissue morphogenesis.

## Discussion

Here, we have shown how optogenetic control can be used to systematically interrogate which features of a developmental signaling pattern are compatible with life. We find that a single pulse of blue light, delivered to both embryonic termini, is sufficient to convert a lethal loss-of-function phenotype to rescue the full *Drosophila* life cycle: embryogenesis, larval development, pupation, adulthood and fecundity.

Our optogenetic rescue result provided two insights into the interpretation of developmental RTK signaling. First, we find that recruiting the catalytic domain of SOS to the plasma membrane recapitulates all the essential developmental functions of Tor receptor tyrosine kinase signaling at the embryonic termini. This complete molecular sufficiency is non-obvious: RTKs activate many intracellular pathways that are bypassed by OptoSOS stimulation (10), some of which have been suggested to play roles in early *Drosophila* embryogenesis (23). Nevertheless, our result is consistent with reports that RTK-driven gene expression (and thus cell fate determination) is mediated primarily by the Ras pathway (24), and that activating Ras pathway mutations are genetic suppressors of Tor partial loss-of-function alleles (25).

Second, our use of simple, all-or-none light inputs suggests that the normally-observed gradient of terminal signaling is dispensable. This statement is supported by three additional observations: (i) all-or-none illumination results in a sharp boundary of OptoSOS membrane translocation (**Figure 2B-C**) (13); (ii) a live-cell Erk biosensor reveals a sharp boundary of light-induced pathway activity compared to the endogenous terminal gradient (**Figure 2B-C**); and (iii) all embryonic positions are uniformly sensitive to SOS stimulation, so uniform illumination over a field induces uniform pathway activity (13). It appears that only two levels of Ras signaling are required – signaling must be “on” above a sufficient threshold at the termini and “off” at interior positions – contrary to a longstanding view of developmental RTK activation (17, 26) and unlike classical morphogens such as the Bicoid transcription factor (27).

Our results suggest that a great deal of the sophisticated information processing in terminal patterning is contained not in the gradient or dynamics of upstream signaling, but rather emerges from downstream transcriptional networks. After all, although simple binary patterns of light are sufficient, we find that distinct developmental events are triggered at vastly different doses (tail structures: 5 min; gastrulation movements: 45 min) and with different spatial requirements (tail structures: no spatial requirement; all other responses: local illumination at the poles). These observations immediately raise an open challenge: we still lack a clear picture of the signal interpretation circuits that respond to different profiles of developmental Erk signaling. Future work combining optogenetic control with high-resolution transcriptional studies (24) to track the localized expression of Erk-responsive terminal genes (e.g. Tll, Hkb) could be crucial for closing this gap.

There is considerable current interest in defining the rules that govern morphogenesis and patterning, both *in vivo* during embryo development and in engineered organoid-based systems. The optogenetic approaches defined here represent a first step toward the delivery light-based programs to specific cells of interest within multicellular tissues. Such a capability may open the door to unprecedented control over developmental processes in both natural and synthetic multicellular systems.

## Author Contributions

H.E.J., S.Y.S. and J.E.T. conceived and designed the project and wrote the manuscript. H.E.J. performed all experiments.

## Acknowledgements

H.E.J. was supported by the NIH Ruth Kirschstein fellowship F32GM119297. This work was also supported by NIH grant DP2EB024247 and NSF CAREER Award 1750663 (to J.E.T.), and NIH grant R01GM086537 (SYS). We also thank Dr. Gary Laevsky and the Molecular Biology Microscopy Core, which is a Nikon Center of Excellence, for microscopy support. Stocks obtained from the Bloomington Drosophila Stock Center (NIH P40OD018537) were used in this study.

