## Supplementary Information for "Optogenetic rescue of a developmental patterning mutant"

### Supplementary Methods

### *Drosophila melanogaster* stocks

### Transgenic UAS-optoSOS flies were produced as described previously (*1*) by fC31-based integration at either CH III 68A4 or CH II 25C6 and are driven maternally using P(mata-GAL-VP16)mat15 (67;15) for all experiments (*2*). For rescue experiments the following LOF lines were used: *trk*^1^, where a premature stop codon prevents production of a functional protein (*3**,* *4*); Torso RNAi (P{TRiP.HMJ22419}attP40) (*5*); and the *tsl*^691^ hypomorphic allele (*6*). The *trk*^1^ and *tsl* lines were recombined with the optoSOS/ 67;15 lines to create the lines used for the rescues. The miniCic-OptoSOS embryos were generated by recombination with OptoSOS CH2 with miniCic (*7*) as well as 67;15 with miniCic, thus producing miniCic-optoSOS/miniCic-67;15/+ mothers.

### Quantification of Erk Activity

Images were obtained of embryos fixed and stained under different pulse frequencies for ppErk (1:100; Cell Signaling). Activity was quantified similar to the method used for gene expression in (*8*) by tracing a contour around the edge of the embryo and normalized based on a co-stained histone-GFP control for each stimulation condition. For quantification of Erk activity during rescue (**Figure 2**), we illuminated individual embryos with an all-or-none light pattern and measured a z-stack in the OptoSOS and miniCic channels. We then computed maximum projections from these z-stacks and plotted the intensity as a function of their distance from the illumination source. For quantifying the endogenous Erk gradient in a wild-type embryo, the maximum Erk activity was measured at the posterior pole. Each curve was then normalized to its minimum and maximum values to obtain estimates of the percentage activity at each position.

### Microscopy

### Cuticles were prepared for imaging as described previously (*1*) and imaged on Nikon Eclipse Ni at 10X objective using dark-field or brightfield microscopy. DIC imaging performed on a Nikon Eclipse Ti spinning-disk confocal microscope. A 740-760nm band-pass filter (Chroma) was placed in the bright-field illumination light path to prevent unwanted optogenetic stimulation during imaging. Patterned optogenetic illumination was performed using a Mightex Polygon digital micromirror device (DMD) using an X-Cite XLED 450-nm blue LED. Spatial illumination of embryos was performed by drawing rectangular DMD patterns to stimulate the anterior pole, posterior pole, or both. The microscope’s XY stage was then cycled between multiple embryos that were approximately aligned to these fixed patterns. Each embryo was stimulated with DMD illumination 600ms every 30s at an intensity of 4.5 mW/cm2, leading to expected activity approximately equal to that of the maximal wild-type level (see Figure S2). Embryos were excluded from subsequent analysis if they were tilted more than 10 degrees from the fixed illumination pattern.

### Assessing and quantifying gastrulation phenotypes

### For experiments classifying gastrulation phenotypes by microscopy, germ band elongation classification was performed based on the raw differential interference contrast (DIC) videos as follows:

### *Terminal loss-of-function-like* – No posterior midgut invagination (PMG), the presence of ectopic folds during gastrulation. Complete absence of germ band extension (GBE).

### *Normal* – The posterior endoderm invaginates and travels along the dorsal surface towards the head. The final posterior-to-anterior length of posterior endoderm migration defines the ‘GBE length’ plotted in Figure S4C.

### *Terminal gain-of-function-like* – Posterior invagination is abnormally large and does not migrate along the dorsal surface, so germband extension is lost.

### The length of GBE was measured at the point which the germ band stops constant forward progress towards the anterior pole. The perimeter of the PMG was measured as the largest contour after the initiation of the PMG invagination. Embryos which exhibited defects prior to NC14, were unfertilized, which were tilted relative to the illumination pattern greater than 10°, or the measured parameter was not visible due to orientation of the embryo were excluded from counts. For Figures S4B-C, embryos which did not receive at least 60 minutes of light between the start of NC10 and 10 minutes before posterior gastrulation movements were also excluded from counts. For Figure S4C, embryos with illumination between 10-30% of EL were included, No GBE ectopic phenotypes were excluded. Embryos were considered WT if found in the range of the 95% confidence interval for OreR controls. For duration experiments the duration which was counted was the light received up until 10 minutes before the start of gastrulation.

### Assessing and quantifying cuticle phenotypes

For cuticle counts, unfertilized /empty eggshells were excluded from all counts. Any visible filzkörper structures were marked as having “tail”. Head structures with intact pharyngeal apparatus were marked as rescued. Abdominal segments were marked as rescued if the cuticle contained 8 intact denticle belts. Hatching rates are based on the number of empty eggshells vs the number of remaining embryos after 30+ hpf.

**Supplementary Figures and Legends**

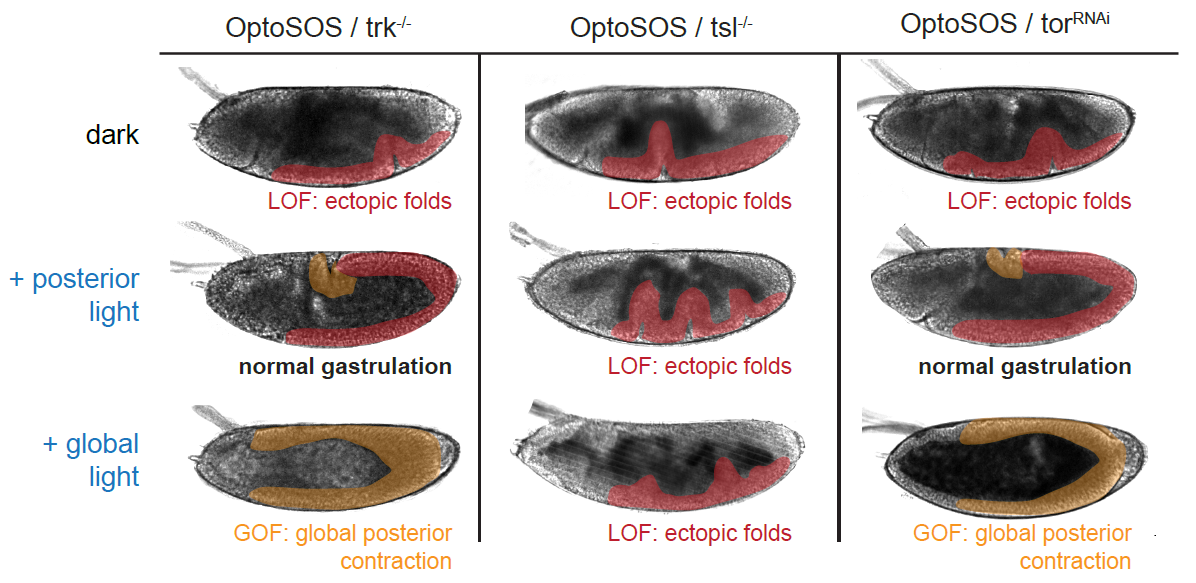

### Figure S1: Characterizing optogenetic control in three Torso signaling pathway mutants. Embryos were analyzed that expressed the OptoSOS system and were derived from mothers harboring loss-of-function Torso pathway alleles: homozygous *trk* (left), homozygous *tsl*, (middle) and 67;15>tor^RNAi^ (right). Each panel shows a representative embryo during gastrulation where the posterior midgut invagination (PMG invagination; yellow highlight) and elongating ventral mesoderm tissue (red highlight) are marked. Top row: all three mutants lead to loss-of-function phenotypes in the dark, including a failure to undergo PMG invagination and the appearance of ectopic folds along the embryo’s ventral surface. Middle row: posterior illumination rescues normal PMG invagination (yellow) and germ band elongation (red) in OptoSOS-*trk* and OptoSOS-tor^RNAi^ embryos. OptoSOS-*tsl* embryos still exhibit loss-of-function gastrulation phenotypes, including a lack of PMG invagination and ectopic folds. Bottom row: global illumination leads to the large-scale formation of ectopic PMG and a massive contraction of the majority of the embryo in OptoSOS-*trk* and OptoSOS-tor^RNAi^ embryos but not in OptoSOS-*tsl* embryos. These data suggest that the *tsl* allele exhibits additional defects in tissue morphogenesis independently of its role in specifying terminal fates.

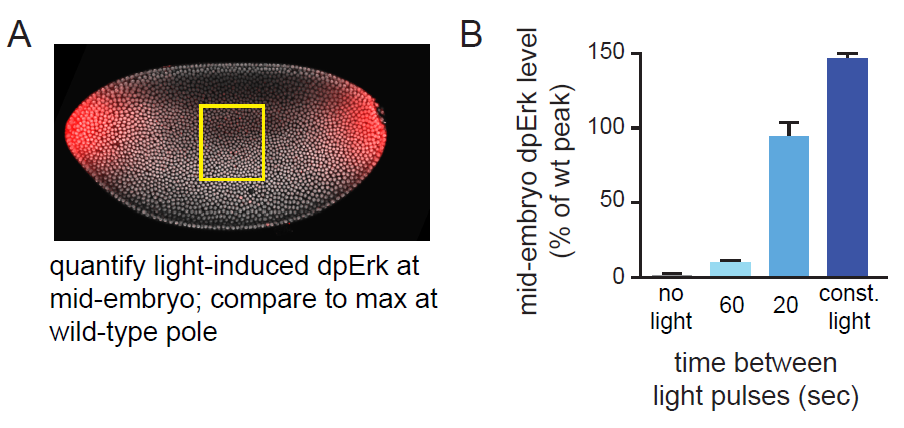

### Figure S2: Titrating Erk phosphorylation using different light stimulus schedules. (A) To quantify Erk activity in response to different light cycles, nuclear cycle 12-14 OptoSOS embryos which had been illuminated for 1-hour prior under different conditions and then fixed and stained for dually phosphorylated Erk (ppErk). The level of ppErk was quantified from an embryonic region far from the termini (middle most quarter of the embryo), so that the only contribution was due to illumination. The maximum level of ppErk at the anterior pole of wild-type embryos was quantified as a relative standard for ppErk intensity. A representative wild-type embryo exhibiting the normal terminal ppErk pattern is shown, reproduced from Figure 1A. (B) Quantification of dpErk levels in embryos stimulated with different pulse cycles: continuous light, a 1 sec pulse every 20 sec, a 1 sec pulse every 60 sec, or constant darkness. Illumination at 1 sec every 20 seconds led to ppErk levels comparable to the maximum level in the wild-type terminal pattern, whereas a 1 sec pulse every 60 sec drove steady-state ppErk levels at ~10% of the wild-type maximum. Error bars: standard deviation across spatial bins.

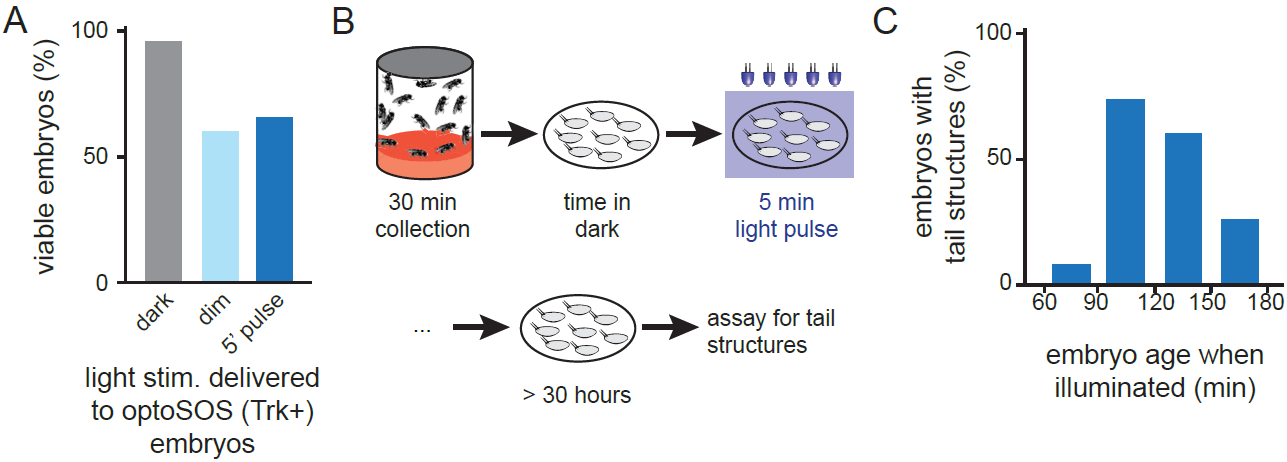

### Figure S3: Programming of tail structures at low Erk levels. (A) Assessing the effects of short-duration and low-intensity illumination on OptoSOS embryos with intact terminal signaling. Embryos were stimulated with identical light stimuli to those used for tail structure rescue in Figure 4 and the fraction of normal embryos was assessed by cuticle preparation. “Dim” light indicates a 1 sec light pulse every 120 sec, expected to lead to Erk activity <10% of maximal levels (see Figure S2). (B) Schematic of experiment to define the time window during which tail structures are specified. Embryos were collected over a 30 min period, incubated in the dark for varying amounts of time, and then globally illuminated with a 5 min light pulse. Cuticle preparations were then used to assess the formation of tail structures at the end of embryogenesis. (C) The fraction of embryos with normal tail structures was defined in each 30 min collection window, with most embryos harboring tails when stimulated in a 1 h window, 90-150 min after fertilization.

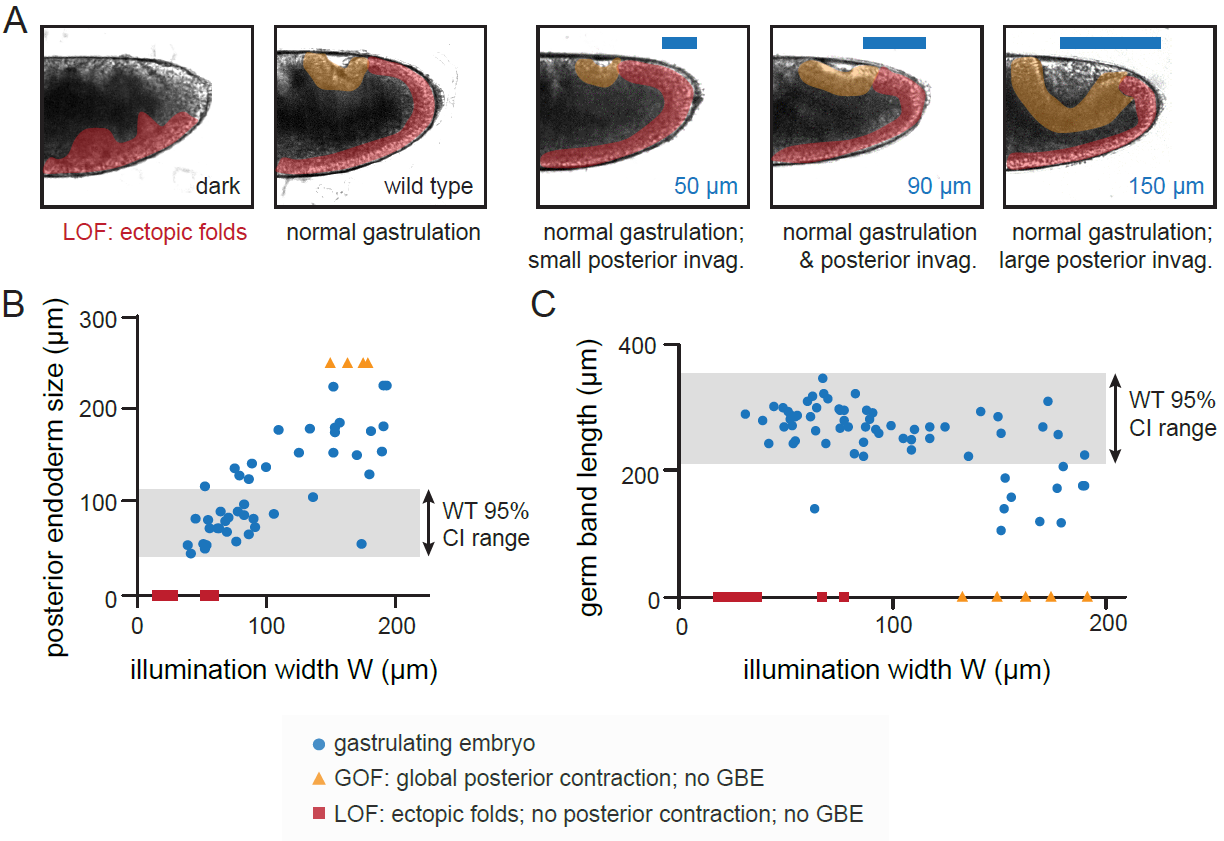

### Figure S4: Tissue morphogenesis is robust to variations in terminal pattern width. (A) Images of gastrulating wild-type embryos and OptoSOS-*trk* embryos that have been stimulated with different illumination widths at the posterior pole. Yellow highlighted regions indicate posterior endoderm invagination, which expands as illumination width is increased. Red highlighted regions indicate the elongating germ band tissue, which buckles in loss-of-function (LOF) embryos. (B-C) Quantification of posterior endoderm perimeter (B) and germ band elongation length (C) as a function of the illumination width from the posterior pole. For some embryos (yellow triangles), posterior contraction was so large as to completely disrupt germ band elongation, a strong gain-of-function (GOF) phenotype. For others (red squares), no posterior contraction occurred, leading to loss-of-function (LOF) failure to extend a germ band at all. For both B-C, the shaded region indicates the normal wild-type size (mean +/- 95% confidence interval), quantified from 50 wild-type embryos.

### Supplementary Table 1: Number of embryos quantified in each experiment. Comma-separated numbers indicate the number of embryos quantified in each experimental condition, from left to right as indicated in the corresponding figure.

| **Figure** | **# of embryos per condition** |
| --- | --- |
| 4A | 204, 90, 68 |
| 4B | 109, 172, 9 |
| 4C | 8, 43, 26, 7, 4, 30 |
| 5B | 10, 44, 34 |
| 5C | 2, 11, 20, 17, 4, 3, 7, 7, 3 (WT=27) |
| S2B | 52, 28, 26, 10 |
| S3A | 88, 21, 117 |
| S3C | 102, 90, 81, 139 |
| S4B | 70 |
| S4C | 82 |

### Supplementary Movie Legends

### Movie S1: Complete rescue of embryogenesis in OptoSOS-*trk* embryo. Time-lapse DIC imaging of a *trk*^1^ optoSOS embryo without (top) and with (bottom) a 90-minute pulse of light at the anterior and posterior poles. Activating light can be seen in blue at the start of the movie.

### Movie S2: Gastrulation in embryo subjected to 150 μm posterior illumination. Time-lapse DIC imaging of an optoSOS-*trk* embryo exposed to a wide posterior illumination pattern (lower panel) compared to a wild-type control (upper panel). An abnormally large posterior invagination is followed by germ band elongation to an extent indistinguishable from a wild-type embryo.
